# A near-tight lower bound on the density of forward sampling schemes

**DOI:** 10.1101/2024.09.06.611668

**Authors:** Bryce Kille, Ragnar Groot Koerkamp, Drake McAdams, Alan Liu, Todd J. Treangen

## Abstract

**Motivation:** Sampling *k*-mers is a ubiquitous task in sequence analysis algorithms. Sampling schemes such as the often-used random minimizer scheme are particularly appealing as they guarantee at least one *k*-mer is selected out of every *w* consecutive *k*-mers. Sampling fewer *k*-mers often leads to an increase in efficiency of downstream methods. Thus, developing schemes that have low density, i.e., have a small proportion of sampled *k*-mers, is an active area of research. After over a decade of consistent efforts in both decreasing the density of practical schemes and increasing the lower bound on the best possible density, there is still a large gap between the two.

**Results:** We prove a near-tight lower bound on the density of forward sampling schemes, a class of schemes that generalizes minimizer schemes. For small *w* and *k*, we observe that our bound is tight when *k* ≡ 1 (mod *w*). For large *w* and *k*, the bound can be approximated by 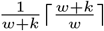. Importantly, our lower bound implies that existing schemes are much closer to achieving optimal density than previously known. For example, with the current default minimap2 HiFi settings *w* = 19 and *k* = 19, we show that the best known scheme for these parameters, the double decycling-set-based minimizer of Pellow et al., is at most 3% denser than optimal, compared to the previous gap of at most 50%. Furthermore, when *k* ≡ 1 (mod *w*) and the alphabet size *σ* goes to ∞, we show that mod-minimizers introduced by Groot Koerkamp and Pibiri achieve optimal density matching our lower bound.

**Availability and implementation:** Minimizer implementations: github.com/RagnarGrootKoerkamp/minimizers ILP and analysis: github.com/treangenlab/sampling-scheme-analysis

## 1 Introduction

For over a decade, *k*-mer sampling schemes have served as a ubiquitous first step in many classes of bioinformatics tasks. By sampling *k*-mers in a way which ensures that two similar sequences will have similar sets of sampled *k*-mers, sampling schemes enable methods to bypass the need to compare entire sequences at the base level and instead allow them to work more efficiently using the sampled *k*-mers.

*Local sampling schemes* satisfy a *window guarantee* that at least one *k*-mer is selected out of every window of *w* consecutive *k*-mers. Most schemes used in practice, such as the random minimizer scheme (Schleimer et al., 2003; Roberts et al., 2004), are *forward schemes* that additionally guarantee that *k*-mers are sampled in the order in which they appear in the original sequence. These properties are particularly appealing since they guarantee that no region is left unsampled.

As the purpose of these schemes is to reduce the computational burden of downstream methods while upholding the window guarantee, the primary goal of most new schemes is to minimize the *density*, i.e., the expected proportion of sampled *k*-mers. Over the past decade, many new schemes have been proposed that obtain significantly lower densities than the original random minimizer scheme.

For example, there are schemes based on *hitting sets* (Orenstein et al., 2016; Marçais et al., 2017, 2018; DeBlasio et al., 2019; Ekim et al., 2020; Pellow et al., 2023; Golan et al., 2024), schemes that focus on sampling positions rather than *k*-mers (Loukides and Pissis, 2021; Loukides et al., 2023), schemes that use an ordering on *t*-mers (*t < k*) to decide which *k*-mer to sample (Zheng et al., 2020; Groot Koerkamp and Pibiri, 2024), and schemes that aim to minimize density on specific input sequences (Zheng et al., 2021b; Hoang et al., 2022). All of these improvements notwithstanding, it is still unknown how close these schemes are to achieving minimum density.

A trivial lower bound on density given by the window guarantee is 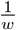, and recently Groot Koerkamp and Pibiri (2024) improved the bound of Marçais et al. (2018) from 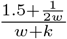 _to 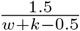_. However, for, there is a sizeable gap between these lower bounds and the density many practical values of *w* and *k* of existing schemes. This raises the question whether schemes with density much closer to 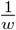 exist, but have not been found yet, or whether existing schemes are already very close to optimal and it is the lower bound that needs improvement. Our new lower bound closes most of the gap, and thus answers this question: Indeed, especially for *k* ≥ *w*, the best existing schemes have near-optimal density in many cases. This allows future research to focus on improving other sampling scheme metrics, such as the *conservation* described in Edgar (2021) and Shaw and Yu (2022).

### 1.1 Contributions

#### Main lower bound theorem

We prove a novel lower bound on the density of forward schemes that is strictly tighter than all previously established lower bounds for all *w, k*, and alphabet size *σ*:

##### Theorem 1.

*Let f be a* (*w, k*)*-forward sampling scheme and M*_*σ*_(*p*) *count the number of aperiodic necklaces of length p over an alphabet of size σ. Then, the density of f is at least*

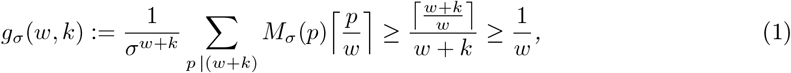

*where the middle inequality is strict for w >* 1.

We prove that this bound can be extended to work for more general classes of sampling schemes, such as the local schemes described by Marçais et al. (2018) and the multi-local schemes described by Kille et al. (2023).

#### Comparison with optimal schemes for small parameters

We show that our lower bound is tight for some small *w, k*, and *σ* by using an integer linear program to construct schemes whose density matches our lower bound. This marks the first time that there is an analytical description of a tight minimum density of any forward scheme. We conjecture that when *k* ≡ 1 (mod *w*), there exist schemes with density matching our lower bound.

#### Comparison with practical schemes for large parameters

To show that our bound is significantly closer to the density achieved by existing schemes compared to previous lower bounds, we replicate the benchmark from Groot Koerkamp and Pibiri (2024) for a selection of *w* and *k* (Fig. 3). For example, with the default minimap2 (Li, 2018) HiFi settings *w* = 19 and *k* = 19, the lower bound goes up from 50% of the density achieved by the double decycling based method to 97% of the achieved density (Table 1).

**Table 1:** Minimum densities achieved by existing sampling schemes for default parameters of frequently-used bioinformatics methods (*σ* = 4). The gap percentage describes the how much larger the lowest achieved density is than the lower bound and is calculated as 100 · 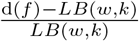 where 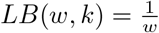 for the old gap and 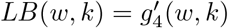 for the new gap. While Groot Koerkamp and Pibiri (2024) showed that 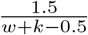 is also a lower bound, 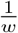 is tighter for all of the parameter choices in the table. For SSHash (Pibiri, 2022), we show parameters used for indexing a single human genome.

| Application | $(w, k)$ | Random<br>$2/(w+1)$ | Best | | Lower bound | | Gap (%) | |
| --- | --- | --- | --- | --- | --- | --- | --- | --- |
| | | | Scheme | Density | $1/w$ | $g'$ | $1/w$ | $g'$ |
| Kraken2 | (5, 31) | 0.333 | Mod-mini | 0.226 | 0.200 | 0.222 | 12.8 | 1.6 |
| SSHash | (12, 20) | 0.154 | Mod-mini | 0.120 | 0.083 | 0.108 | 43.9 | 10.9 |
| minimap2, hifi | (19, 19) | 0.100 | dbl decycling | 0.079 | 0.053 | 0.077 | 50.1 | 2.7 |

#### Analysis of the mod-minimizer

Finally, our new lower bound implies that the mod-minimizer scheme (Groot Koerkamp and Pibiri, 2024) is optimal when *k* ≡ 1 (mod *w*) and *σ* is large. Indeed, for the ASCII alphabet (*σ* = 256), the mod-minimizer scheme density is consistently within 1% of the lower bound when *k* ≡ 1 (mod *w*) (Fig. 4 in Supplement D).

## 2 Background

### Notation

We begin by defining some necessary notation, as well as definitions of mathematical concepts that will be used throughout the work. We use [*n*] to refer to the set {0, 1, …, *n* − 1}. The alphabet is denoted by Σ and has size *σ* := |Σ|, with *σ* = 4 for DNA. The expression *a* | *b* indicates that *a* divides *b*. The summation Σ_*a*|*b*_ is over all positive divisors *a* of *b*. We use *a* mod *m* for the remainder (in [*m*]) of *a* after dividing by *m* and we use *a* ≡ *b* (mod *m*) to indicate that *a* and *b* have the same remainder modulo *m*. Given a string *W, W* [*i, j*) refers to the substring of *W* containing the characters at 0-based positions *i* up to *j* − 1 inclusive. For two strings *X* and *Y, XY* represents the concatenation of *X* and *Y*.

### Classes of sampling schemes

There are multiple established classes of sampling schemes. We begin by drawing a distinction between schemes with and without a *window guarantee* that guarantees that at least one every *w k*-mers is sampled. While schemes without a window guarantee, such as fracminhash (Irber et al., 2022), are often efficient to compute, the lack of a guarantee on the distance between sampled *k*-mers makes them ineffective or inefficient for certain tasks such as indexing and alignment. Indeed, we only consider schemes with a window guarantee:

#### Definition 1.

*A* (*w, k*)-local scheme *with window guarantee w and k-mer size k on an alphabet* Σ *corresponds to a sampling function f* : Σ^*w*+*k*−1^ → [*w*].

In other words, given a window of *w* + *k −* 1 characters (*w* consecutive *k*-mers), the output of the sampling function *f* (*W*) is an integer in [*w*] which represents the index of the sampled *k*-mer in *W*. Recently, Kille et al. (2023) proposed a generalization of (*w, k*)-local schemes which samples at least *s k*-mers out of every *w* instead of at least 1 and we extend our results to these more general schemes in Supplement A.

Local schemes have no restrictions on which of the *w k*-mers can be selected for each window, but *forward schemes* are a subset of local schemes that enforce the restriction that they never select a *k*-mer which occurs before a previously selected *k*-mer.

#### Definition 2.

*A* (*w, k*)*-local scheme is also* (*w, k*)-forward *if for all strings W* ∈ Σ ^*w*+*k*^ *representing two adjacent windows*,

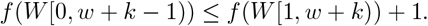

#### Definition 3.

*The* density d (*f*) *of as ampling scheme f is defined as the expected proportion of sampled positions from an infinite, uniformly random string*.

For a further background on types of sampling schemes, we refer to Shaw and Yu (2022), Zheng et al. (2023), Groot Koerkamp and Pibiri (2024), and Ndiaye et al. (2024).

### De Bruijn graphs

Let *B*_*n,σ*_ = (*V, E*) denote the complete De Bruijn graph of order *n*, which has as vertices all strings of length *n, V* = Σ^*n*^, and edges between vertices that overlap in *n −* 1 positions, *E* = {(*X, X*[1, *n*)*c*) | *X* ∈ *V, c* ∈ Σ}. When *σ* is clear from the context or irrelevant for a particular discussion, it is omitted. It is worth noting that the vertices of *B*_*n*+1_ correspond to edges of *B*_*n*_.

For each string *s* of length *n*, the *n* rotations of *s* induce a *pure cycle* in *B*_*n*_ consisting of (up to) *n* vertices cyclically connected by edges. Note that when *s* is repetitive, e.g., a single repeated character or some other repeated string, the length of the cycle will be a divisor of *n*. These pure cycles are also called *necklaces*. The set of necklaces of length *n* corresponds to a partitioning of the vertices of *B*_*n*_ into a vertex-disjoint set of pure cycles. We use C_*n*_ to refer to this set of pure cycles of *B*_*n*_, and for *c* ∈ C_*n*_, we write |*c*| for the number of vertices in the cycle.

When a string of length *n* has *n* unique rotations, the corresponding necklace is said to be *aperiodic*. The total number of necklaces and the number of aperiodic necklaces of length *n* are given by Moreau (1872) (and see also Riordan (1957)) as respectively

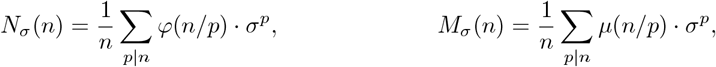

where *φ*(*p*) is Euler’s totient function that counts the number of integers in [*p*] coprime to *p*. The formula *M*_*σ*_(*n*) counting *aperiodic* necklaces follows from the formula for *N*_*σ*_(*n*) via *Möbius inversion* (Möbius, 1832), where *µ* is the *Möbius function* defined to be 0 if *n* is divisible by a square (*>* 1) and *µ*(*n*) = (−1)^*q*^ otherwise, where *q* is the number of prime factors of *n*.

### Charged contexts

The *context* of a window of length *w* + *k −* 1 in a sequence is the set of preceding windows that influences whether the current window samples a new position.

For a local scheme to select a new position, none of the previous *w* − 1 windows may have selected the same *k*-mer as the current window. As a result, the context for local schemes consists of 2*w* + *k* − 2 characters: the current window of *w k*-mers as well as the *w* − 1 windows preceding the current window. For a forward scheme, however, as soon as a window samples a different position than the preceding window, this position must be a new position. Thus, one needs only to consider the context of two consecutive windows of *w k*-mers, for a total of *w* + *k* characters.

When a sampling scheme selects a new position for the last window in a context, the context is *charged*. Marçais et al. (2017) showed that the density of a scheme *f* can be defined as the proportion of contexts which are charged. In the case of forward schemes, each edge in *B*_*w*+*k*−1_ represents a context, and the charged contexts are the edges (*u, v*) for which *f* (*u*)≠ *f* (*v*) + 1.

### Universal hitting sets

In 2021, Zheng et al. (2021a) related the density of forward and local schemes to the concept of universal hitting sets (UHS). A (*w, 𝓁*)-UHS is defined as a set of *𝓁*-mers *U* such that *any* sequence of *w* adjacent *𝓁*-mers must contain at least one *𝓁*-mer from *U*. Theorem 1 of Zheng et al. (2021a) showed that when *k* = 1, one can use the minimum size of a (*w, 𝓁*=*w* + *k*)-UHS to bound the density of (*w, k*=1)-forward schemes, and the minimum size of a (*w, 𝓁*=2*w* + *k* − 2)-UHS to bound the density of a (*w, k*=1)-local schemes.

## 3 Theoretical results

In this section, we prove our main result: an improved lower bound on the density of forward sampling schemes. We first generalize some existing theorems to arbitrary *w* and *k* (Sections 3.1 and 3.2), after which our main theorem follows in Section 3.3.

### 3.1 A lower bound on the size of a (*w, 𝓁*)-UHS

We begin by considering a (*w*=2, *𝓁*)-UHS. A (2, *𝓁*)-UHS is equivalent to a vertex cover in *B*_*𝓁*_, i.e., a subset of vertices such that each edge in *B*_*𝓁*_ is adjacent to at least one vertex in the subset. Lichiardopol (2006) used the fact that for every cycle *C*, at least ⌈|*C* |*/*2⌉ of its vertices must be in a vertex cover, and obtained a lower bound on the size of a vertex cover by partitioning *B*_*𝓁*_ into its pure cycles. We naturally extend this argument to obtain a lower bound on the cardinality of a (*w, 𝓁*)-UHS for any *w* ≥ 2.

#### Proposition 4.

*Let M*_*σ*_(*p*) *count the number of aperiodic necklaces of length p. For any* (*w, 𝓁*)*-UHS*

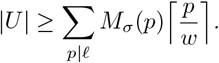

*Proof*. The pure cycles of C_*𝓁*_ partition the vertices of *B*_*𝓁*_. For any simple cycle of size *p* in *B*_*𝓁*_, a (*w, 𝓁*)-UHS must contain at least ⌈*p/w*⌉ *𝓁*-mers. As there is a one-to-one correspondence between the pure cycles of length *p* | *𝓁* in *B*_*𝓁*_ and the *M*_*σ*_(*p*) aperiodic necklaces of length *p*, we have

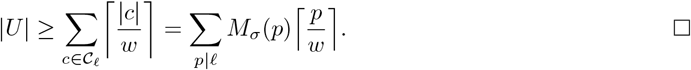

Fig. 1b provides a depiction of a minimum (2, 4)-UHS as well as the pure-cycle partitioning of *B*_4_ on a binary alphabet. Notably, the pure cycle (0011, 0110, 1100, 1001) has 3 vertices in the UHS, even though the lower bound given by Proposition 4 only requires it have 2. This is an example where the lower bound is not tight.

**Figure 1:**
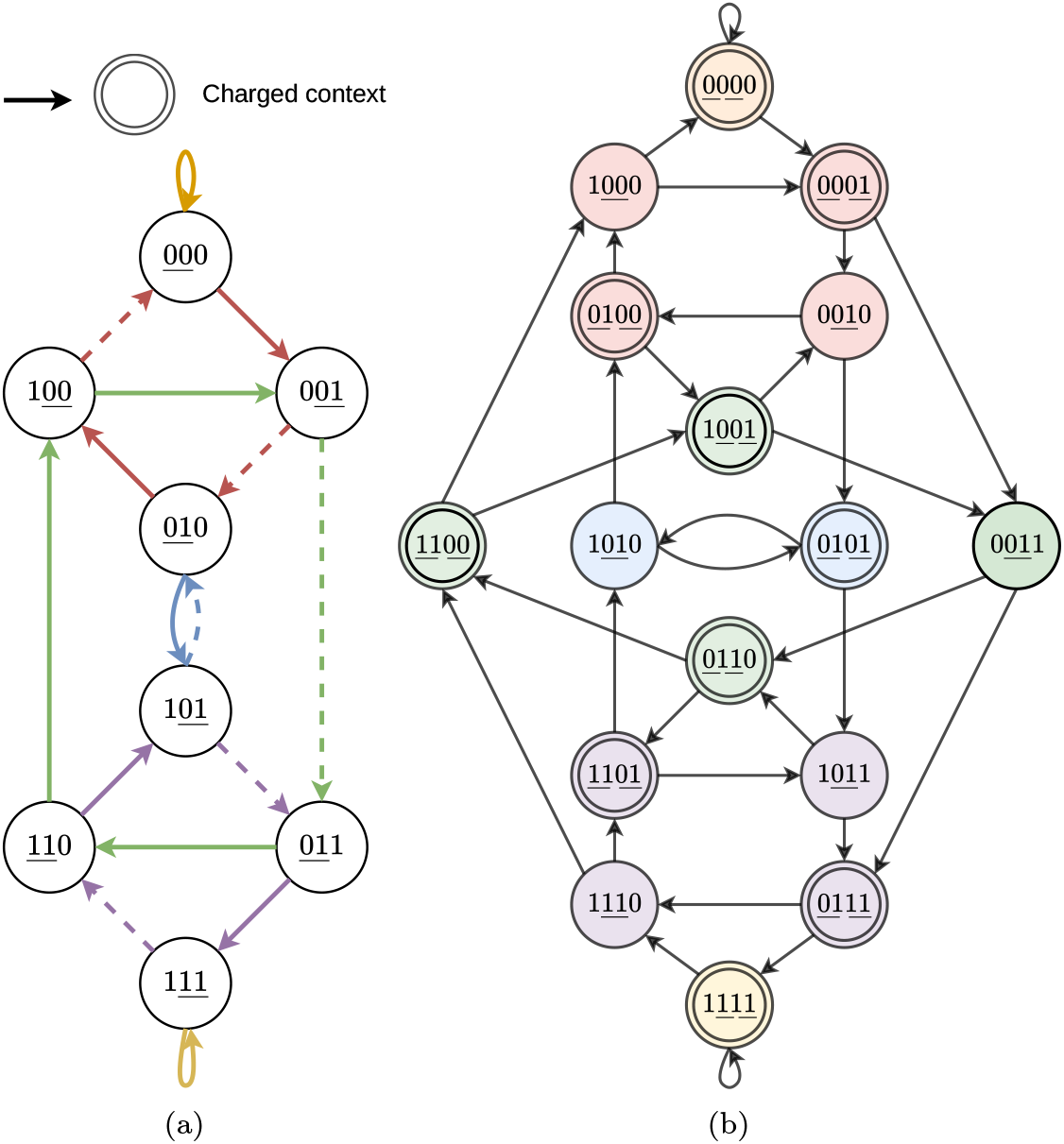
(a) A De Bruijn graph *B*_3_ corresponding to a minimum density (*w*=2, *k*=2)-forward scheme. The underlined characters in each vertex represent the 2-mer that is selected for that window. The solid edges represent the charged contexts and the edge colors represent the pure cycles in *B*_4_ (not in *B*_3_ itself). For characters *c*_*i*_, each edge 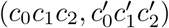 in *B*_3_ corresponds to the vertex 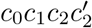 in *B*_4_. (b) The corresponding (*w*=2, *𝓁*=4)-UHS in *B*_4_. The vertices are partitioned by color, representing the pure-cycles. The 2-mer(s) selected in each context are underlined. The vertices with a double border represent the charged edges in *B*_3_ in (a) and the corresponding (2, 4)-UHS. Each pure cycle *c* has at least ⌈|*c* |*/w*⌉ vertices in the UHS.

For certain values of *w* and *𝓁*, such as when *𝓁* is prime or *w* = 2 and *𝓁* is odd, Proposition 4 can be simplified to remove the summation and ceil function (Supplement B).

Proposition 4 is the core of the proof of Theorem 1 and already has the right structure. The remainder of this section translates this result on universal hitting sets to a result on the density of sampling schemes.

### 3.2 A connection between sampling scheme density and UHS size

Zheng et al. (2021a, Theorem 1) show a connection between universal hitting sets and the density of sampling schemes when *k* = 1. We naturally extend their result to *k* ≥ 1 for both local schemes (Lemma 5) and forward schemes (Corollary 6).

#### Lemma 5.

*Let f be a* (*w, k*)*-local scheme, and let C*_*f*_ *be its corresponding set of charged contexts defined as the set of strings W of length* 2*w* + *k* − 2 *for which the last window W* [*w* − 1, 2*w* + *k* − 2) *selects a position w* − 1 + *f* (*W* [*w* − 1, 2*w* + *k* − 1)) *not selected by any previous window:*

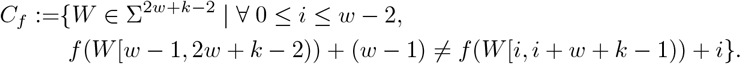

*Then, C*_*f*_ *is a* (*w*, 2*w* + *k* − 2)*-UHS*.

*Proof*. For the sake of a contradiction, suppose there is a walk of length *w* in the De Bruijn graph of order (2*w* + *k* − 2), say (*W*_0_, …, *W*_*w*−1_), that avoids *C*_*f*_. Let *S* be the *spelling* of the walk, i.e., the sequence of length 3*w* + *k* − 3 such that *S*[*i, i* + 2*w* + *k* − 2) = *W*_*i*_. Since *W*_*w*−1_ ∈*/ C*_*f*_ and *S* contains *W*_*w*−1_, this implies that on the last (*w* + *k* − 1)-mer of *W*_*w*−1_ (i.e., *S*[2*w* − 2, 3*w* + *k* − 3)), *f* selects an index *j* ≥ 2*w* − 2 in *S* which has already been picked.

Since 0 ≤ *f* (·) ≤ *w* − 1 and *j* ≥ 2*w* − 2, the first window that selects position *j* must begin at an index *m* ≥ *w* − 1. Therefore, the context *W*_*m*−*w*+1_ = *S*[*m* − (*w* − 1), *m* + *w* + *k* − 1) is charged, as *f* selects a previously unselected position when applied to its last (*w* + *k* − 1)-mer. By definition, *W*_*m*−*w*+1_ ∈ *C*_*f*_, contradicting our supposition and therefore *C*_*f*_ is a (*w*, 2*w* + *k* − 2)-UHS.

Identically, one can consider contexts for a (*w, k*)-forward scheme *f*, which requires only verifying that the selection for a window of length *w* + *k −* 1 is distinct from the selection for the previous window. Therefore, the length of a context for forward *f* is only *w* + *k*. As above, every *w* contexts must have at least one charged context, leading to the following conclusion:

#### Corollary 6.

*If f is a* (*w, k*)*-forward scheme and C*_*f*_ *is its corresponding set of charged contexts, defined as C*_*f*_ = {*W* ∈ Σ^*w*+*k*^ | *f* (*W* [0, *w* + *k* − 1))≠ *f* (*W* [1, *w* + *k*)) + 1}, *then C*_*f*_ *is a* (*w, w* + *k*)*-UHS*.

As all contexts of a particular length *𝓁* are equally likely to occur in an infinite, uniform random string, the proportion of charged contexts corresponds to the density of the sampling scheme (Marçais et al., 2017), i.e. d(*f*) = |*C*_*f*_| */σ*^*𝓁*^, where *𝓁* = *w* + *k* for forward schemes and *𝓁* = 2*w* + *k* − 2 for local schemes. An example of the charged contexts of a (2, 2)-forward scheme and the corresponding UHS is depicted in Fig. 1.

### 3.3 Lower bounds on local and forward scheme density

We are now ready to state and prove our main theorem.

#### Theorem 1.

*Let f be a* (*w, k*)*-forward sampling scheme and M*_*σ*_(*p*) *count the number of aperiodic necklaces of length p over an alphabet of size σ. Then, the density of f is at least*

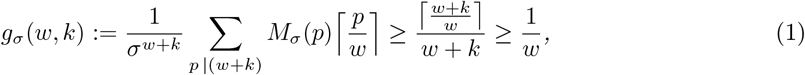

*where the middle inequality is strict for w >* 1.

*Proof*. Due to Corollary 6 and Marçais et al. (2017), we can see that a (*w, k*)-forward sampling scheme of density d(*f*) implies a (*w, 𝓁*=*w* + *k*)-UHS of size *σ*^*w*+*k*^ · d(*f*). By Proposition 4, this implies that every forward sampling scheme has a density of at least *g*_*σ*_(*w, k*), and hence d(*f*) ≥ *g*_*σ*_(*w, k*) follows.

For any *p* that divides *w* + *k*, we have 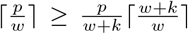, with strict inequality when *p* = 1 and *w >* 1. Substituting this in *g*_*σ*_(*w, k*), the middle inequality follows directly using the identity Σ_*p*|*w*+*k*_ *p* · *M*_*σ*_(*p*) = *σ*^*w*+*k*^ that counts the number of strings of length *w* + *k* partitioned by their shortest period.

The last inequality follows directly from 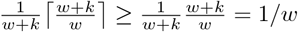.

As we will see in Section 4, *g*_*σ*_(*w, k*) is a tight bound for many small cases. Since its formula is somewhat unwieldy 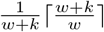 can be used as an approximation that quickly approaches *g*_*σ*_(*w, k*) (Fig. 2). Simple arithmetic shows that both *g*_*σ*_(*w, k*) and 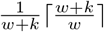 improve the previous lower bound of 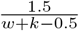 of Groot Koerkamp and Pibiri (2024).

**Figure 2:**
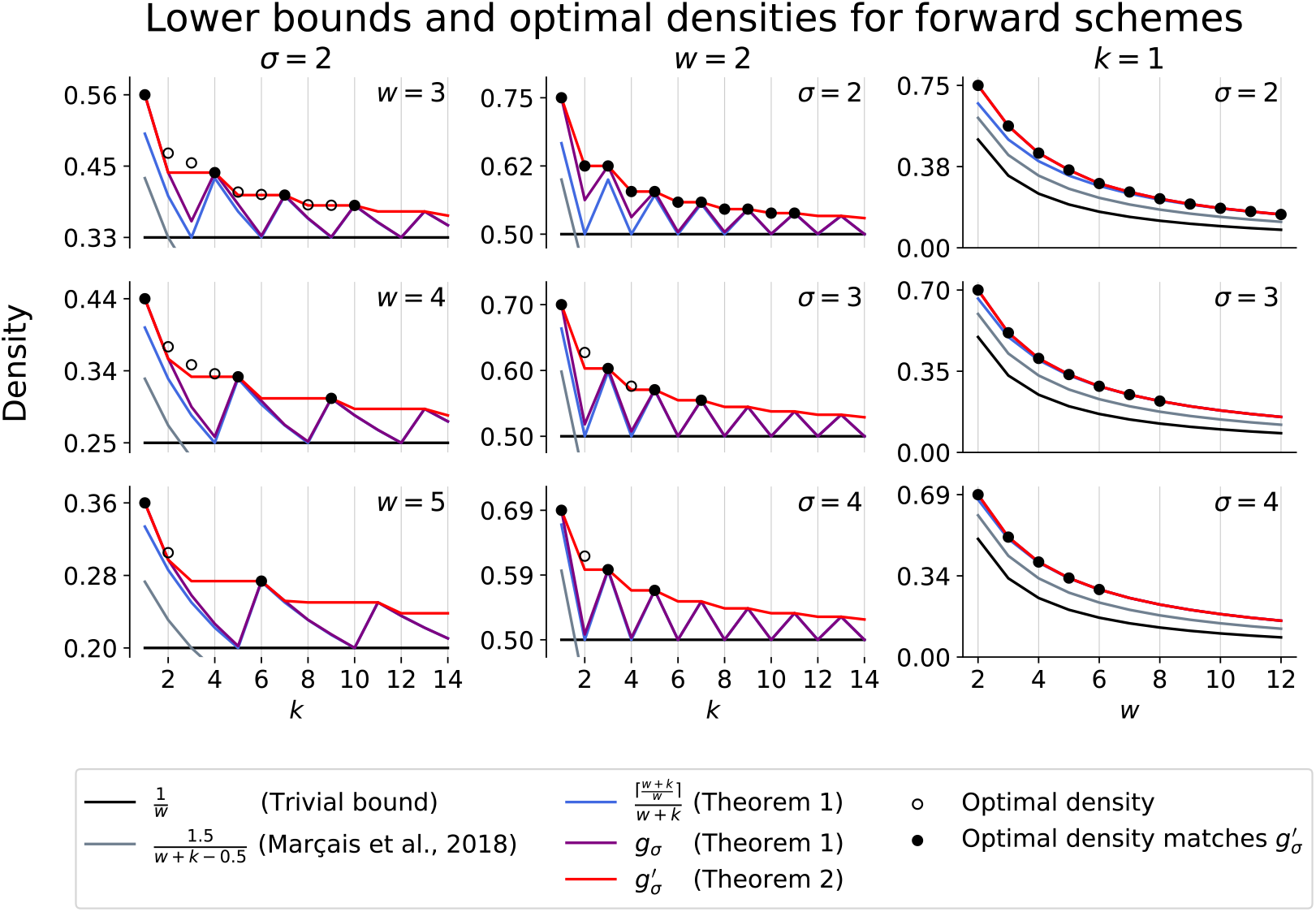
Comparison of forward scheme lower bounds and optimal densities for small *w, k*, and *σ*. Optimal densities were obtained via the ILP and are plotted as black circles that are solid when the optimal density matches our lower bound, 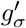, and hollow otherwise. Each column corresponds to a parameter being fixed to the lowest non-trivial value, i.e., *σ* = 2 in the first column, *w* = 2 in the second column, and *k* = 1 in the third column. Note that the x-axis in the third column corresponds to *w*, not *k*.

**Figure 3:**
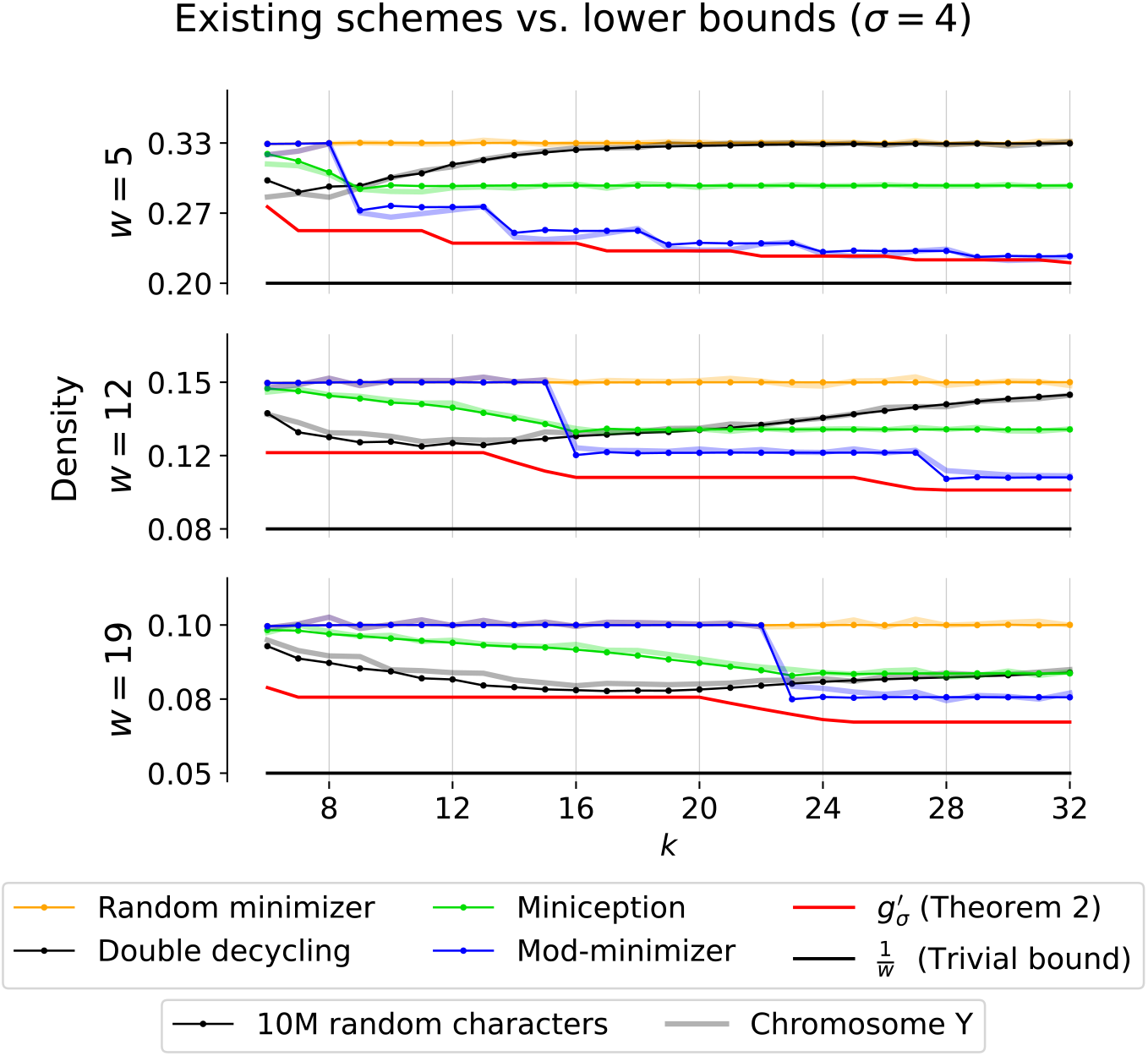
Comparison of existing schemes to lower bounds with practical parameters. Densities are calculated by applying each scheme to a random sequence of 10 million characters over an alphabet of size *σ* = 4 (dotted lines) and are compared with the corresponding proportion of sampled *k*-mers on the human Y chromosome (Rhie et al., 2023) (soft lines). The mod-minimizer uses parameter *r* = 4, and miniception uses parameter max(4, *k − w*). The window sizes 5 and 19 are the default window sizes for Kraken2 (Wood et al., 2019) and minimap2 (`-ax hifi`) (Li, 2018), respectively. For SSHash, *w* = 12 was the window size used when indexing the human genome (Pibiri, 2022).

**Figure 4:**
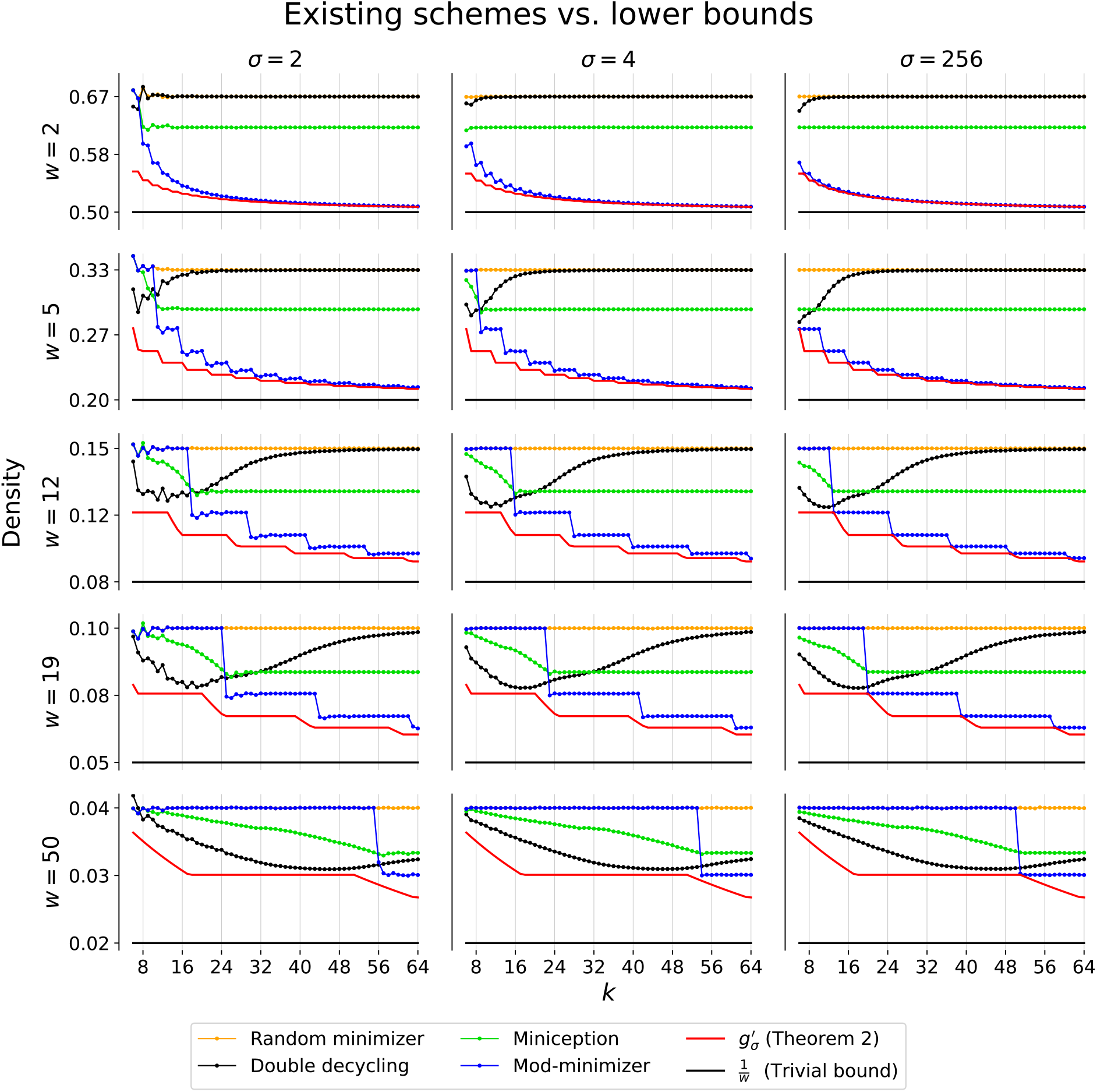
Comparison of existing schemes to lower bounds. Densities were calculated by applying each scheme to a random sequence of 10 million characters and are plotted as solid dotted lines. The mod-minimizer uses parameter *r* = 6 for *σ* = 2, *r* = 4 for *σ* = 4, and *r* = 1 for *σ* = 256. Miniception uses parameter max(*r, k* − *w*). Lower bounds are plotted as solid lines.

Given a (*w, k*)-local scheme *f*_*k*_, we can construct a (*w, k* ≥ *k*)-local scheme *f*_*k*_*′* of the same density by ignoring the last *k*^*′*^ *− k* characters in each window, i.e. *f*_*k*_*′* (*W*) = *f*_*k*_(*W* [0..(*w* + *k*)). This directly implies d(*f*_*k*_) = d(*f*_*k*_*′*) (Zheng et al., 2021a). It follows that the minimum density of a (*w, k*)-local or forward scheme is monotonically decreasing as *k* increases. However, as can be seen in Fig. 2, *g*_*σ*_(*w, k*) is not a monotonically decreasing function. The local maxima appear to be at *k* ≡ 1 (mod *w*), which motivates the following improved lower bound.

#### Theorem 2.

*For any* (*w, k*)*-forward scheme f, an improved lower bound g*^*′*^ *is given by*

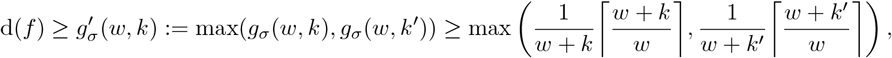

*where k*^*′*^ *is the smallest integer* ≥ *k such that k*^*′*^ ≡ 1 (mod *w*).

#### Remark 7.

*Similar to Theorem 1, Lemma 5 implies that any* (*w, k*)*-local scheme f has density at least* d(*f*) ≥ *g*_*σ*_(*w, w* + *k* − 2). *As this bound is in terms of g*_*σ*_, *the improved bound in Theorem 2 can be applied to local schemes as well, i*.*e*., *for any* (*w, k*)*-local scheme f, an improved lower bound is given by*

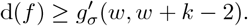

## 4 Empirical tightness of our bounds

Here, we compare our bounds *g*_*σ*_ and 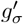 to existing lower bounds. Further, we show how tight these bounds are for small *w, k*, and *σ* by searching for optimal schemes via an integer linear programming (ILP) formulation. We also show how close existing sampling scheme densities are to 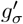 for practical choices of *w, k*, and *σ*. Finally, we show when the recently described mod-minimizer scheme (Groot Koerkamp and Pibiri, 2024) achieves optimal density as *σ* → ∞.

### ILP description

We use an ILP to search for minimum density forward sampling schemes. In short, we use a single integer variable *x*_*W*_ ∈ [*w*] for every window *W* of length *w* + *k* − 1 (corresponding to a vertex in *B*_*w*+*k*−1_) that indicates the position of the chosen *k*-mer, and a single boolean variable *y*_(*W,W ′*)_ for each edge in *B*_*w*+*k*−1_ that indicates whether the corresponding context is charged. On each edge, we require that the scheme be forward. The objective function is to minimize the number of charged edges. To reduce the search space, we add an additional constraint corresponding to our lower bound *g*_*σ*_ by requiring that for each pure cycle of length |*c*| in *B*_*w*+*k*_, at least ⌈|*c*|*/w*⌉ of the corresponding edges in *B*_*w*+*k*−1_ are charged. Further details, including the ILP formulation for local schemes, can be found in Supplement C.

### Comparison against optimal schemes for small *k*

We used Gurobi (Gurobi Optimization, LLC, 2024) to solve the ILP for all combinations of *w, k*, and *σ* such that 1 ≤ *w* ≤ 12, 1 ≤ *k* ≤ 12, and 2 ≤ *σ* ≤ 4 for both forward and local schemes and limited the runtime for each instance to 12 hours on 128 threads. All results are reported in Table 2 in Supplement D. While the additional constraint on pure cycles corresponding to *g*_*σ*_ significantly sped up the search, for most large *w, k*, and *σ*, the ILP failed to terminate with an optimal solution in the allotted time. As a result, we restrict most of our analysis to the following three cases: fixed alphabet size *σ* = 2, fixed window size *w* = 2, and fixed *k*-mer size *k* = 1 (Fig. 2).

**Table 2:**
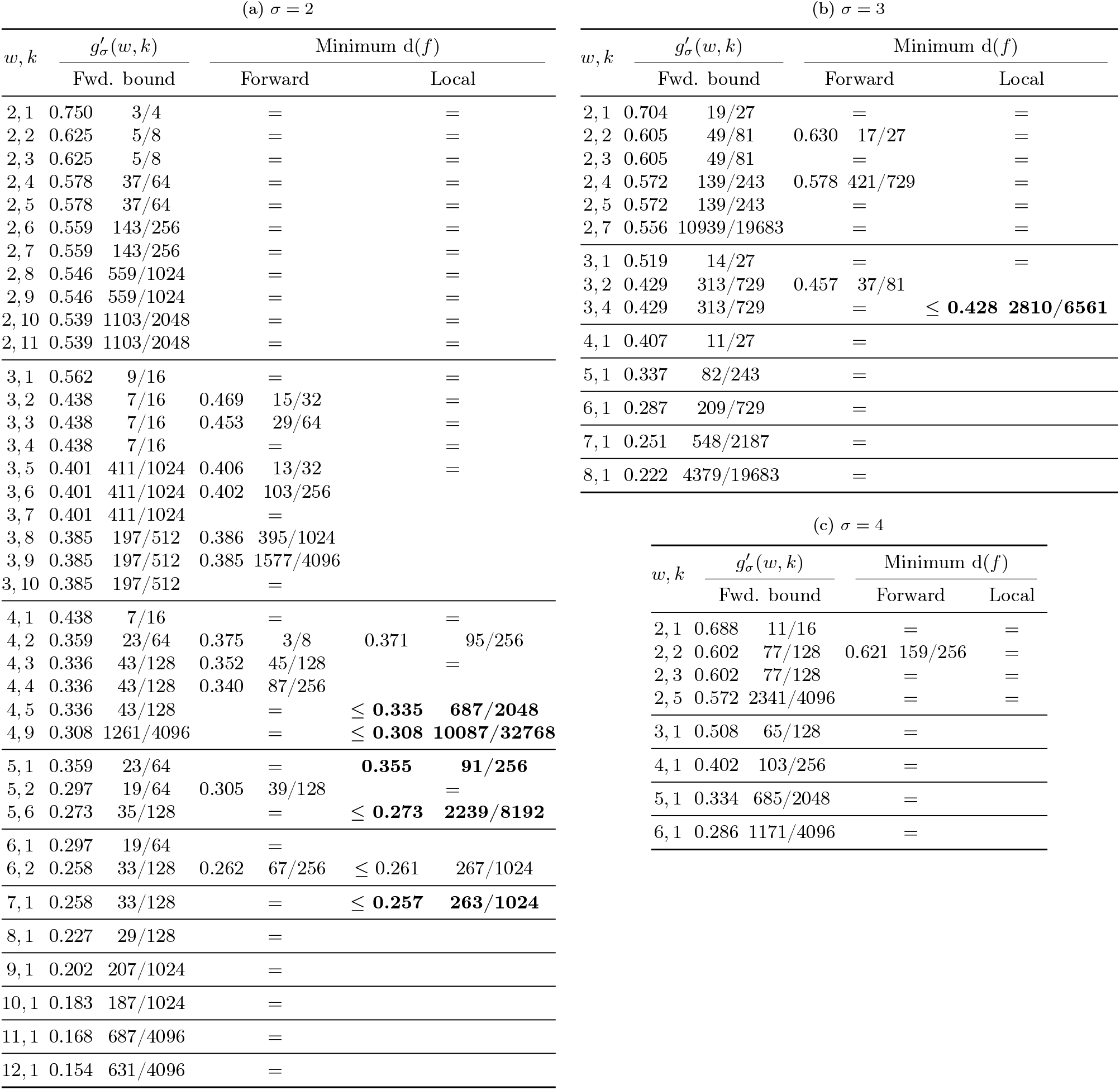
Our 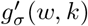 lower bound on the density of forward schemes, and minimum densities of forward and local obtained by the ILPs described in Supplement C. Entries with an equals sign (=) correspond to parameters where the minimum density is equal to the preceding column. Entries with the less-than-or-equal-to sign (≤) correspond to parameters where the local ILP identified a solution with lower density than the optimal forward scheme but timed out before determining whether the identified solution was optimal. Bold entries indicate cases where the local scheme is better than the *g*^*′*^(*w, k*) bound for forward schemes. For *w* = 2, all local schemes are forward by definition and therefore the minimum (2, *k*)-local scheme density is equal to the minimum (2, *k*)-forward scheme density. Empty cells correspond to parameters where the local ILP timed out before it was able identify a local scheme with lower density than the optimal forward scheme or prove that no such scheme exists. In these cases, it is unknown whether or not there exists a local scheme with lower density than the optimal forward scheme.

For all (*w, k, σ*) where *k* ≡ 1 (mod *w*) (including when *k* = 1), the minimum density exactly matches our lower bound *g*_*σ*_(*w, k*). Additionally, when *σ* = 2 and *w* = 2, the minimum density was equal to 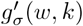.

### Comparison against existing schemes for large *k*

Using a sequence of 10 million random characters over alphabet size *σ* = 4, we approximated the density of recent sampling schemes using the benchmarking implementation from Groot Koerkamp and Pibiri (2024). To compare each density to the particular proportion of selected *k*-mers on a genomic sequence, we also ran all sampling schemes on the human Y chromosome (Rhie et al., 2023) after removing all non-ACTG characters. The densities of the best performing methods, Miniception (Zheng et al., 2020), double decycling-set-based minimizers (Pellow et al., 2023), and mod-minimizers (Groot Koerkamp and Pibiri, 2024) are plotted in Fig. 3 along with random minimizers and lower bounds.

The ratio between the minimum achieved densities and lower bounds for a selection of (*w, k*) pairs used by existing *k*-mer based methods are presented in Table 1. Additional results for *σ* ∈ {2, 256} and *w* ∈ {2, 50} are provided in Fig. 4 in Supplement D.

### The mod-minimizer has optimal density for large *σ* when *w* ≡ *k* (mod 1)

When *w* and *k* are constant and *σ* → ∞, the probability of duplicate characters in a window goes to 0. This implies that we can use *t* = 1 for the mod-minimizer. When *k* ≡ *t* = 1 (mod *w*), the density of the mod-minimizer (Theorem 10 of Groot Koerkamp and Pibiri (2024)) is given by

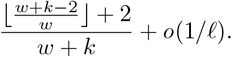

The *o*(1*/𝓁*) term only accounts for duplicate *t*-mers, and hence disappears when *σ* → ∞. We get

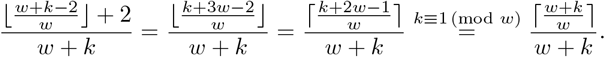

Thus, the mod-minimizer has density equal to the lower bound provided by Theorem 1 when *σ* goes to ∞ and *w* and *k* ≡ 1 (mod *w*) are fixed.

In practice, for *σ* = 256 the mod-minimizer scheme is within 1% from optimal when *k* ≡ 1 (mod *w*) (Fig. 4 in Supplement D). When *σ* = 4 (Fig. 3), a *t >* 1 must be used, causing the density plot to “shift right” compared to the lower bound. Because of that, the mod-minimizer does not quite match the lower bound for practical values of *σ*.

## 5 Discussion

### 5.1 Conjecture on when our lower bound is tight

Analytically, it is clear that 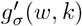 is much larger than 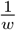. In all cases, 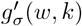 is nearly tight, if not completely. In particular, our bound is tight for all 40 tested parameter sets where *k* ≡ 1 (mod *w*), leading us to our conjecture:

#### Conjecture 1.

*For any w and k satisfying k* ≡ 1 (mod *w*), *there exists a* (*w, k*)*-forward sampling scheme f such that* d(*f*) = *g*_*σ*_(*w, k*).

While the minimum size of a decycling set, i.e., a (*w* =∞, *𝓁*)-UHS, is well-known to be *N*_*σ*_(*𝓁*) (Mykkeltveit, 1972), very little is known about the minimum size of a (*w, 𝓁*)-UHS for finite *w*. In addition to providing the minimum density of a (*w, k*)-forward scheme for *k* ≡ 1 (mod *w*), proving Conjecture 1 would also determine the minimum size of a (*w, 𝓁*=*w* + *k*)-UHS when *k* ≡ 1 (mod *w*).

### 5.2 Existing schemes are nearly optimal when *k* ≥ *w* or *σ* is large

A natural investigation which follows our proposed lower bound is to determine the gap between 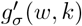 and current forward scheme densities. Previously, the gap between known densities and lower bound was rather large, making it unclear how much more the density could be reduced.

In Table 1, we observe that existing schemes are already within 11% from the optimal density for practical values of *w* and *k* across different applications, and in many cases are even within 3% of the optimal density. In Fig. 3, we see that this difference holds not just for the specific (*w, k*) in Table 1, but for most *k* ≥ *w*. This is much more informative than the previous lower bound of 1*/w*, which implied that most current schemes are at most 50% denser than optimal for many of the parameters in Fig. 3.

For alphabets much larger than DNA (*σ* = 4), such as the ASCII alphabet (*σ* = 256), we observe that when *k* ≡ 1 (mod *w*), the mod-minimizer scheme recently proposed by Groot Koerkamp and Pibiri (2024) is at most 1% denser than optimal and furthermore, we show that it is optimal as *σ* → ∞. This makes the mod-minimizer scheme the first practical scheme for which there exist finite parameters *k* and *w* for which it is close to optimal.

### 5.3 Tightening the bound for small *k*

Our new bound for forward schemes always improves over 1*/w* and appears tight when *k* ≡ 1 (mod *w*). This leads to an increasingly close bound for *k*≢ 1 (mod *w*) as *k* increases, but leaves a large gap when 1 *< k < w*. A better understanding of these small cases will be necessary to obtain a tight lower bound for all *w* and *k*. Looking at Fig. 3 and Fig. 4 in Supplement D, one might conjecture that the double decyling-set-based methods Pellow et al. (2023) are near-optimal, but subsequent work such as the greedy minimizer (Golan et al., 2024) has shown better schemes are possible. From Fig. 2, we already know that our lower bound is not always tight, so this leaves the question:

**Open problem 1**. *How close can practical sampling schemes get to the density given by our lower bound?*

### 5.4 Extending the bound to local schemes

For local schemes, though, our bound appears much less tight. We identified 8 sets of (*w, k, σ*) where local schemes can obtain lower densities than their forward counterparts. In all cases, however, the difference between the local and forward densities was minuscule, with the largest difference of being found for (*w* = 4, *k* = 2, *σ* = 2) where the density decreased from 0.375 to 0.371 (Table 2 in Supplement D). Nevertheless, for some parameters, local schemes are able to achieve densities lower than our 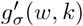 lower bound for forward schemes. Given the trend observed in Table 2, we arrive at our final open problem:

**Open problem 2**. *How much lower can the density of a* (*w, k*)*-local scheme be compared to a* (*w, k*)*-forward scheme?*

## Acknowledgements

We thank Giulio Ermanno Pibiri, Nicolae Sapoval, and our reviewers for their suggestions which helped improve the manuscript.

## A (*s, w, k*)-multi-local schemes

Given a set *A*, we define 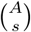 as the set of all subsets of *A* of cardinality *s*.

### Definition 8.

*A* (*s, w, k*)-multi-local scheme *corresponds to a sampling function* 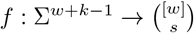, *where s is the sketch size, w is the window guarantee, k is the k-mer size, and σ* = |Σ| *is the alphabet size*.

A quick substitution reveals that (*s, w, k*)-multi-local schemes are equivalent to (*w, k*)-local schemes when *s* = 1. These schemes enable a stricter window guarantee by ensuring that at least *s k*-mers are selected for every window of *w k*-mers. However, it was recently shown that this generalization can also be used to outline a more relaxed window guarantee (Kille et al., 2023).

Consider the case where instead of desiring at least one sampled *k*-mer for every *w k*-mers, the requirement is to have *s k*-mers sampled for every *sw k*-mers. This latter goal can be accomplished by a (*w, k*)-local scheme, but as was shown in Kille et al. (2023), a much lower density can be obtained with an (*s, sw, k*)-multi-local scheme.

Here, we will show how the bounds in the main text can be extended to yield a lower bound for (*s, w, k*)-multi-local schemes.

A (*s, w, 𝓁*)-multi-UHS is defined as a mapping *α* : Σ^*𝓁*^ → [*s* + 1] that assigns weights to vertices in *B*_*𝓁*_ such that any sequence of *w* adjacent *𝓁*-mers must have a combined weight of at least *s*. Again, when *s* = 1, this corresponds to the (*w, 𝓁*)-UHS described in the main text. We now provide an extension to Proposition 4 which provides a lower bound on the size of a (*s, w, 𝓁*)-multi-UHS.

### Proposition 9.

*For any* (*s, w, 𝓁*)*-multi-UHS α*,

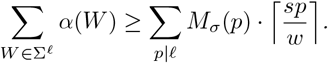

*Proof*. We first show that for any simple cycle of size *p* in *B*_*𝓁*_, the combined weight of the cycle must be at least 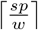. Let *W*_0_, …, *W*_*p*−1_ be the vertices in a cycle of length *p*. Consider the walk of length *w* along the cycle starting at *W*_0_. We have that Σ _*i*∈[*w*]_ *α*(*W*_*i*_) ≥ *s* where indices are taken modulo *p*. This same inequality holds for all *p* unique paths of length *w* along the cycle and summing over all of them yields 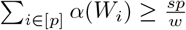. Finally, since Σ_*i*∈[*p*]_ *αα*(*W*_*i*_) must be an integer and each pure cycle in C_*𝓁*_ of length *p* corresponds to an aperiodic necklace of length *p*, of which there are *M*_*σ*_(*p*), we arrive at the result:

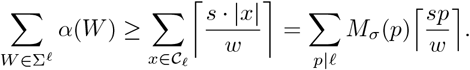

The multi-local schemes also require that we generalize our definition of a charged context. As a multi-local scheme can select multiple new *k*-mers in a single window, we change our binary notion of a charged context to a mapping of weights to contexts, where a context *W* has weight *α*(*W*) if *α*(*W*) previously unsampled positions are selected in the final window of length *w* + *k* − 1 in *W*. Similar to local schemes, a multi-local scheme’s necessary context is the current window of *w k*-mers as well as the previous *w* − 1 windows, leading to a context size of 2*w* + *k* − 2.

**Lemma 10**. *If f is a* (*s, w, k*)*-multi-local scheme and α* : Σ^2*w*+*k*−2^ → [*s* + 1] *is the corresponding mapping of weights to contexts, defined as α*(*W*) = |*A\ B*|, *where A consists of all positions in the context selected by the final window and B consists of all positions in the context selected in all previous windows, i*.*e*.

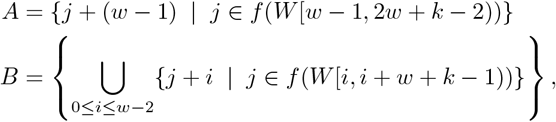

*then αα is a* (*s, w*, 2*w* + *k* − 2)*-multi-UHS*.

*Proof*. Our proof follows a similar structure to that of Lemma 5. Let us show that the total weight of any path of length *w* in *B*_2*w*+*k*−2_ is at least *s*. Let a sequence of *w* consecutive contexts of length 2*w* + *k* − 2 be given as (*W*_0_, …, *W*_*w*−1_). Take *S* to be the sequence of length 3*w* + *k* − 3 such that *S*[*i*, 2*w* + *k* − 2 + *i*) = *W*_*i*_. Then we have that *f* on the last (*w* + *k* − 1)-mer of *W*_*w*−1_ (which is *S*[2*w* − 2, 3*w* + *k* − 3)) selects *s* distinct indices *i*_1_, *i*_2_, …, *i*_*s*_ in *S* where for each 1 ≤ *p* ≤ *s* we have *i*_*p*_ ≥ 2*w* − 2. Suppose *n* of these indices are selected by *f* on any previous (*w* + *k* − 1)-mer of *S*, indexed 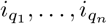, while the other *s* −*n i*_*p*_ are not (meaning *α*(*W*_*w*−1_) = *s* −*n*). Clearly, if *n* = 0, we are done. Therefore, assume instead *n >* 0.

Recall that for all 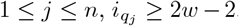. Since 0 ≤ *f* (·) ≤ *w* − 1, we have that for each *j*, the first (*w* + *k* − 1)-mer *S*[*m*_*j*_, *m*_*j*_ + *w* + *k* − 1) in *S* such that *f* picks the index 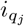 satisfies *w* − 1 ≤ *m*_*j*_ ≤ 2*w* − 2. Then for each *j*, the choice of *m*_*j*_ yields that 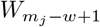 selects a new location 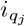 when *f* is applied to its last (*w* + *k* − 1)-mer. This means that 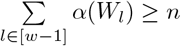, since the sum accounts for every context of the form 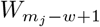, and hence every chosen index 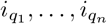. As *α*(*W*_*w*−1_) = *s* − *n*, we have that the sum of the weights of the *w* contexts *W*_0_, …, *W*_*w*−1_ is at least *s*.

Tying these results together using the same line of reasoning as we did in the main text for local schemes, we arrive at a lower bound for multi-local schemes:

### Corollary 11.

*If f is a* (*s, w, k*)*-multi-local scheme, then*

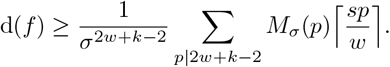

Let us consider the use case of multi-local schemes described earlier, i.e., where the requirement is to have *s k*-mers sampled from every *sw k*-mers. We showed in the main text that this can be accomplished by a (*w, k*)-forward scheme with density at least *g*_*σ*_(*w, k*), or a (*w, k*)-local scheme with density at least *g*_*σ*_(*w, w* + *k* − 2). With our new bound, we can see that the requirement can be achieved with a (*s, sw, k*)-multi-local scheme with density at least *g*_*σ*_(*w, k* + (2*s* − 1)*w* − 2). In other words, the lower bound for local schemes is the same pattern as the forward bound, but “shifted left” by *w* − 2, and the bound for multi-local is the same as the local bound, but again shifted left by (2*s* − 2)*w*.

## B An alternative form of *g*_*σ*_(*w, k*)

When all divisors of *𝓁* apart from 1 have the same remainder modulo *w*, we can simplify *g*_*σ*_(*w, k*).

### Corollary 12.

*Let N*_*σ*_(*𝓁*) *denote the number of cycles in the pure cycle partitioning of B*_*𝓁*_. *Let w, 𝓁 be a pair of integers such that w does not divide 𝓁 and for all divisors d* | *𝓁 excluding the unit divisor* 1, *d* ≡ *z* (mod *w*). *Then for any* (*w, 𝓁*)*-UHS U*, 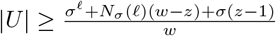.

*Proof*. There are *σ* singleton cycles in a De Bruijn graph on an alphabet of *σ* characters, and each of these must be included in any hitting set *U*. For all remaining cycles, we have that ⌈|*c*|*/w*⌉ = (|*c*| + *w* − *z*)*/w*.

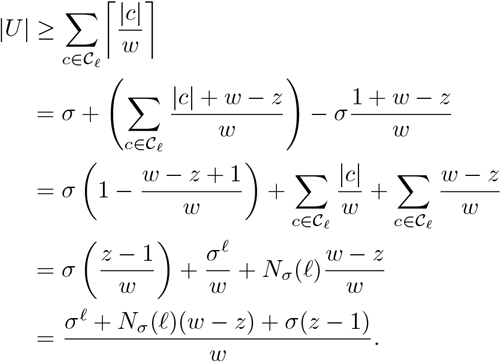

While the formula in Corollary 12 still includes a summation over divisors due to its use of *N*_*σ*_, it no longer involves any ceiling calculations. Furthermore, it shows that *g*_*σ*_(*w, k*) can be written as the following when *w* and *k* satisfy the constraints of Corollary 12

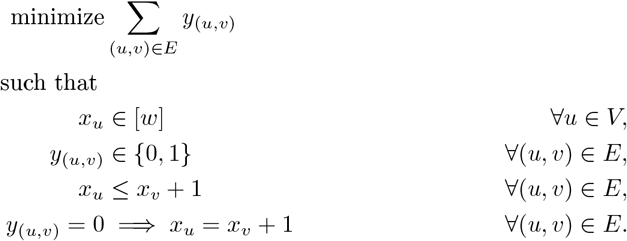

This form, compared to the general form of *g*_*σ*_(*w, k*), provides a more interpretable characterization of the gap between *g*_*σ*_ and 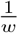.

## C ILP Model

### Forward ILP definition

We used Gurobi (Gurobi Optimization, LLC, 2024) to implement our ILP. While the basic model described below is sufficient, we made multiple improvements which enabled identifying solutions for larger *σ, w* and *k*. We use *x*_*u*_ to model *f* (*u*), i.e. the index of the *k*-mer selected in the window *u* ∈ Σ^*w*+*k*−1^. The *y*_*u,v*_ variables represent the charge of a context of two adjacent windows *u* and *v*.

Given a De Bruijn graph *B*_*w*+*k*−1_ = (*V, E*), the windows correspond to the vertices of the graph, and the contexts correspond to the edges. An edge (*u, v*) is not charged if *x*_*u*_ = *x*_*v*_ + 1. For a forward scheme, *x*_*u*_ ≤ *x*_*v*_ + 1 for every edge (*u, v*). Therefore, we can define an ILP which minimizes the density of a (*w, k*)-forward scheme as follows:

### Local ILP definition

While we could define an ILP for local schemes similarly, the resulting model is inefficient due to the reliance on intermediate variables. Instead, we leverage the fact that the expected density on a random string (Marçais et al., 2017) is the same as the density of a (*w, k*)-local scheme on a circular De Bruijn sequence of order 2*w* + *k* − 2.

Let *S* be circular De Bruijn sequence of order 2*w* + *k* − 2, i.e. *S* is a circular sequence of length *L* = *σ*^2*w*+*k*−2^ which contains every (2*w* + *k* − 2)-mer exactly once. As *S* is circular, we note that when *i > j*, the substring *S*[*i*..*j*) corresponds to *S*[*i*..*L*)*S*[0..*j*), We use *x*_*W*_ to represent the value of *f* (*W*) and *y*_*i*_ to be a binary variable which corresponds to whether or not position *i* was sampled. We define an ILP which minimizes the density of a (*w, k*)-local scheme as follows:

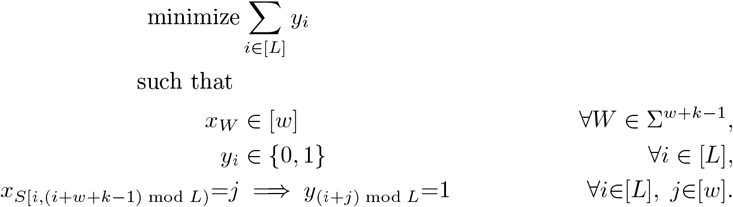

### Improvements to the base forward ILP

First, we added the additional constraint that each pure cycle *c* in *B*_*w*+*k*_ must have at least ⌈|*c* | */w*⌉ nodes which correspond to charged edges in *B*_*w*+*k*−1_. This additional constraint helps substantially when *k* ≡ 1 (mod *w*). In cases where *k* ≢ 1 (mod *w*), we add corresponding constraints for a subset of simple cycles in *B*_*w*+*k*_ that correspond to pure cycles in higher or lower order De Bruijn graphs. While this process adds many more constraints to the model, it narrows the search space substantially and also increases the objective of the ILP linear relaxation, leading to a smaller (and sometimes nonexistent) integrality gap.

As adding this constraint requires the model to have variables representing the charge of a context, it was more efficient to directly use the context charges as the objective variable as opposed to using selected positions in a De Bruijn sequence (as we do in the local ILP). For the local ILP, the *g*_*σ*_ bound is relatively loose and therefore the aforementioned cycle constraints do not help much. Furthermore, modeling context charges requires more intermediate variables in the local ILP, hence we directly used sampled positions in a De Bruijn sequence as the objective for the local ILP.

If *g*_*σ*_(*w, k*) is tight for some *σ, w, k*, then the objective value of the linear relaxation of the ILP is the same as the integer objective value. As a result, the objective bound is fixed from the start. We leveraged this fact by telling the optimizer to focus on identifying integral solutions as opposed to decreasing the gap between the objective bound and objective value in cases where we suspect that *g*_*σ*_(*w, k*) is tight. This was done through the heuristics parameter.

Finally, we used previously computed solutions to provide starting points for the ILP optimization. Let *f* be a (*w, k*)-local scheme. We constructed a (*w* + 1, *k*)-local scheme or a (*w, k* + 1)-local scheme *f* ^*′*^ through ignoring the last character in an input window *W* of length *w* + *k*, i.e. *f* ^*′*^(*W*) = *f* (*W* [0, *w* + *k*)). This is similar to the “naive extensions” described in Marçais et al. (2018). If we had a minimum density (*w* − 1, *k*)-forward scheme or a (*w, k* − 1)-forward scheme, we seeded the optimizer with the “extended” sampling function which corresponded to whichever one of the precursors schemes yielded the lower density.

We ran our ILP model on a server with 128 threads and 128GB of RAM on all combinations of 2 ≤ *w ≤* 12, 1 ≤ *k ≤* 12, and 2 ≤ *σ ≤* 4. For each set of parameters, we limited the runtime to 12 hours. The ILP identified optimal forward schemes for 60 different sets of parameter and optimal local schemes for 11 different sets of parameters.

## D Additional figures

See Fig. 4 and Table 2.

